# Network analysis reveals how lipids and other cofactors influence membrane protein allostery

**DOI:** 10.1101/2020.07.06.187484

**Authors:** Annie M. Westerlund, Oliver Fleetwood, Sergio Perez-Conesa, Lucie Delemotte

## Abstract

Many membrane proteins are modulated by external stimuli, such as small molecule binding or change in pH, transmembrane voltage or temperature. This modulation typically occurs at sites that are structurally distant from the functional site. Revealing the communication, known as allostery, between these two sites is key to understanding the mechanistic details of these proteins. Residue interaction networks of isolated proteins are commonly used to this end. Membrane proteins, however, are embedded in a lipid bilayer which may contribute to allosteric communication. The fast diffusion of lipids hinders direct use of standard residue interaction networks. Here, we present an extension which includes cofactors such as lipids and small molecules in the network. The novel framework is applied to three membrane proteins: a voltage-gated ion channel (KCNQ1), a G-protein coupled receptor (GPCR - *β*2 adrenergic receptor) and a pH-gated ion channel (KcsA). Through systematic analysis of the obtained networks and their components, we demonstrate the importance of lipids for membrane protein allostery. Finally, we reveal how small molecules may stabilize different protein states by allosterically coupling and decoupling the protein from the membrane.

## I. INTRODUCTION

Proteins are molecular machines which function via complex networks of interacting residues. A perturbation at a specific site in a protein may travel across the protein to modulate a distal site. The specific chain of residues that propagate this information to the distal site constitutes an *allosteric pathway* [1]. Allostery is the major functional contributor in membrane proteins such as receptors and ion channels [2–5]. G-protein coupled receptors (GPCRs), for example, bind specific ligands at an extracellular binding site. These binding events yield conformational changes at an intracellular binding site, altering the receptor’s active/inactive equilibrium [6]. Voltage-gated ion channels, on the other hand, sense and respond to membrane potentials in domains which are located far away from the pore gate. Activation of these domains are allosterically communicated to the pore to trigger channel opening [7–11]. Understanding protein allostery and its governing components may therefore reveal the mechanistic details of these proteins.

All-atom molecular dynamics (MD) simulations provide atomistic details of how protein residues interact with each other, as well as with their surroundings. MD simulations thus contain explicit information about protein allostery. However, the large number of atoms in physiologically relevant systems prevents direct visual identification of residues that are especially important for communicating allostery. This requires computational methods able to synthesize and interpret the raw data. Residue interaction networks of isolated proteins have previously been used to reveal protein allostery [11–15]. Membrane proteins, however, are surrounded by lipids and may be modulated by small molecules, both of which play an integral role in communicating allostery between different protein domains [2, 16, 17]. The original framework therefore has to be extended to reveal the allosteric role of cofactors, such as lipids, ligands or highly retained water molecules.

The difficulty of including cofactors in these networks relates to their redundancy, and the network topology. A network consists of two components: nodes and connection between nodes (edges), which are static. Cofactors, however, constantly move and may exchange positions. To understand the role of lipids, for example, the main goal is to reveal whether a lipid bound in a specific location relative to the protein is allosterically important, rather than a specific lipid molecule which may diffuse away. The redundancy of cofactors therefore has to be translated to the nodes of the static network.

Here, we extend the framework of standard residue interaction networks to include cofactors such as lipids and ligands. In the Theory section, we explain how to assign cofactor residues to network nodes by solving a combinatorial optimization problem in each trajectory frame, i.e. minimizing the distances between cofactor residues and network nodes. This framework is applied in the Results section to MD simulations of three membrane proteins with different functions and topologies: a voltage-gated ion channel, a GPCR, and a pH-gated ion channel. By applying current flow analysis [18, 19] to these networks, we reveal their allosteric fingerprints. Finally, we access the allosteric role of lipids, and show how this role may be modulated by ligands. The method can easily be applied to other systems. A jupyter notebook tutorial is publicly available on our lab github [20].

## II. THEORY

### A. Constructing a residue interaction network

An ordinary residue interaction network is built by grouping the atoms of each protein residue to a node and forming edges between nodes of interacting residues. The weight of an edge represents the extent of residue interaction. Practically, the network is here obtained from the elementwise product of a contact map, *C*, and a matrix representing the correlation of node movement, *M*,

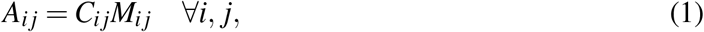

where *A* is the network adjacency matrix. Fig. 1 shows a flowchart of how the network is built using MD simulation data. We use a continuous contact map and positional mutual information to model spatial proximity and correlation of residue movements, respectively (see details below) [21]. The analysis is performed on non-hydrogen atoms of the protein residues, ligand and lipids. MDtraj [22] is used as backbone to perform analysis.

**FIG. 1.**
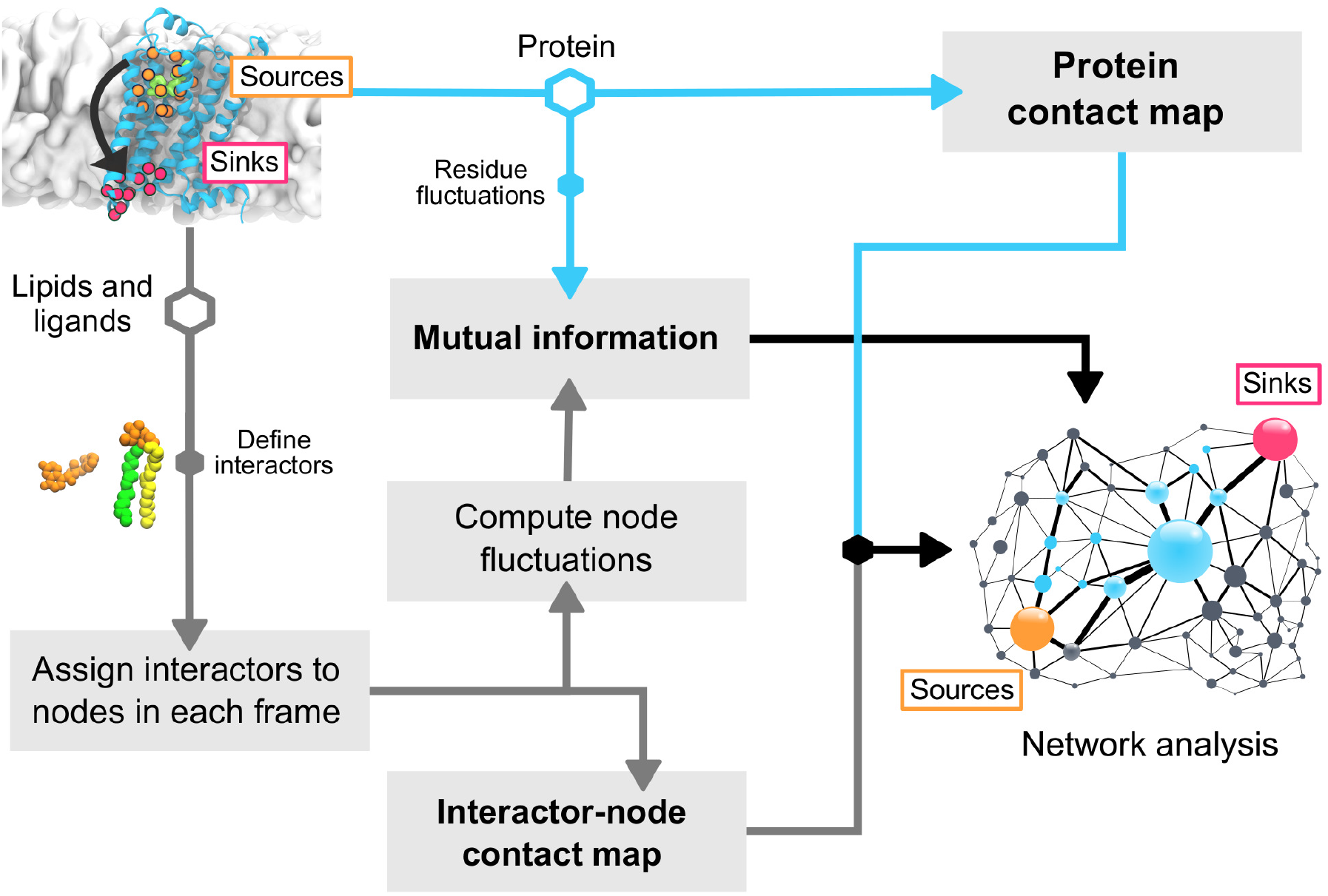
Steps carried out to build a residue interaction network which includes lipids and ligands as well as the protein. Starting from a system with a protein embedded in a lipid membrane and bound ligand, the protein is first isolated. Each protein residue is assigned a node in the isolated protein network and the contact map and mutual information for all protein residue pairs are calculated (blue arrows) The lipids and ligands are divided into “interactors”. These interactors are assigned network nodes depending on their positions relative to the initial frame. The nodes are used to compute the interactor-protein contact map (gray arrows), and the fluctuations are used to compute mutual information between node movements. The contact maps are merged. The full contact map and mutual information make up the final network (black arrows). Once the network is built, we can extract allosteric pathways between source (orange) and sink (red) nodes employing network analysis.

### B. Assigning lipids and ligands to nodes

The network can be extended to represent cofactors as additional nodes. Similar to the protein, we define *interactors* by grouping cofactor atoms into domains. These interactors will correspond to interactor-nodes in the network. Here, we describe a ligand with one interactor while lipids are divided into three interactors; one corresponding to the head group and two to the hydrophobic tails, Fig. 1, S1.

A first solvation shell lipid may exchange position with a second shell lipid. In such case, the protein environment is essentially unaltered, while the individual lipid environments change abruptly. We therefore keep the network nodes static by permuting the lipid interactors and reassigning interactors to nodes in each frame. To do this, the trajectory is first aligned on the protein C_*α*_ atoms. Nodes are assigned to interactors in the first frame of the trajectory. Next, we permute the interactor-nodes in each trajectory frame such that the network remains similar to the initial frame. In practice, for a specific frame, we wish to assign each interactor node to a node in the initial frame network. This is done by minimizing all centroid distances between pairs of the specific and initial frame nodes [23]. We solve this optimization problem with a linear assignment problem solver (the python package LAPJV) [24–26]. The centroid distances are computed with the minimum image convention using an orthorhombic box periodic boundary condition.

### C. Continuous contact map

Continuous contact maps yield more stable networks than binary ones [14]. A switching-function, *K*(*d*) introduced in [21] is used. Given the minimum distance *d*_*ij*_(*n*) between atoms of nodes *i* and *j* in frame *n*,

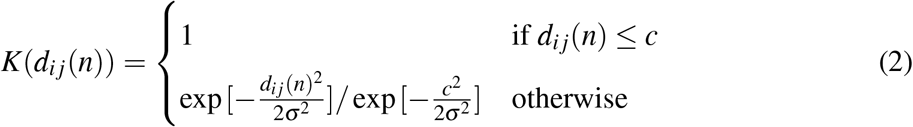

Distances below *c* are given a weight of 1, while weights of larger distances are continuously decreased. The cutoff *c* = 4.5Å is suitable to define contacts between heavy atoms [27] and is commonly used in binary contact maps. We therefore used *c* = 0.45 nm, and *K*(*d*_*cut*_) = 10^*−*5^, where *d*_*cut*_ = 0.8 nm, leading to 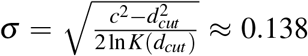. The chosen *d*_*cut*_ allows for small deviations from the hard cutoff, *c*, to be included as weaker interactions. The final contact map is averaged over frames,

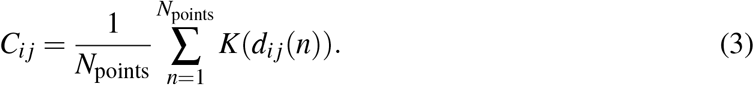

The minimum image convention is used to compute distances involving interactor nodes [22]. This makes the network periodically connected, allowing allosteric pathways to cross simulation box boundaries.

### D. Mutual information

Correlation of residue and interactor node movements are modeled with positional mutual information (MI) of node centroids [21, 28]. This measures correlation of fluctuation distances around an equilibrium position by characterizing the amount of information that can be inferred about a specific node position knowing the position of another node. Given the fluctuation density, *ρ*_*i*_(*x*), where *x* is the distance to the equilibrium position, the entropy of residue *i* is calculated as

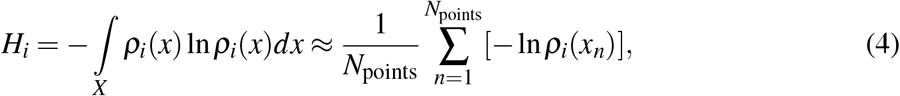

and the MI between nodes *i* and *j* is obtained with

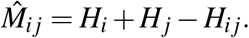

To improve accuracy of entropy estimation, we sample *N*_points_ new data points from the density, *ρ*_*i j*_(*x*), 10 times, followed by repeating entropy and MI calculations and averaging over datasets.

Each density is represented by a Gaussian mixture model (GMM) [29]; a linear combination of Gaussian basis functions. Because Gaussians are continuous, the density estimation is more stable where the data is sparse and does not introduce unnecessary approximations compared to discrete methods such as histogramming [30, 31]. Given the number of Gaussians, GMM parameters (amplitudes, means and covariances) are estimated with expectation-maximization [29]. Increasing the number of Gaussians arbitrarily increases the likelihood. To choose a non-overfitting but detailed model, we use the Bayesian information criterion (BIC) [32], a criterion often used for mixture models [33]. BIC adds a penalty to the likelihood, which increases with model complexity. The GMM with the smallest BIC is selected. Ultimately, BIC is prior-free with respect to GMM parameters, and assumes a flat prior over models such that the most likely model is chosen [33]. For each density, we vary the number of Gaussians between 1 and 4.

Because non-protein interactors may cross the simulation box boundaries, their node equilibrium positions are calculated by first replicating the node frame coordinates to all periodic images. The replication creates clusters of points. One cluster corresponds to the fluctuations around one image equilibrium, which is given by the centroid of this cluster. We then identify a full cluster using a cutoff around the node’s first frame coordinate in the original box. The cutoff to identify the full cluster is taken as half the smallest box side. Although this cutoff may seem conservative for connecting all points within the same cluster, it effectively avoids connecting points that are part of different clusters. The equilibrium position is then defined as the cluster centroid.

## III. INFERRING ALLOSTERY WITH INFORMATION FLOW ANALYSIS

*Information flow*, often called *current flow* after the original analogy to electrical networks [18, 19], measures the net flow of information between a set of source (*S*_0_) and sink (*S*_1_) nodes. Allostery within a protein is communicated via multiple pathways [1]. As opposed to shortest path betweenness [34], current flow betweenness takes all pathways between source and sink nodes into account (details in Supplementary Material) [19]. The net flow across edges naturally removes uninteresting regions in which ineffective pathways might move back and forth before reaching the sink. Because of this, explicit pathways may sometimes be difficult to infer directly. Nonetheless, the result highlights nodes that carry effective pathways, making current flow betweenness well-suited to extract allosterically important residues and interactors.

Current flow betweenness was first introduced to biomolecular simulations in the context of a soluble enzyme [14]. A homomeric-averaged variant was recently presented and experimentally validated on KCNQ1 [21]. The current flow betweenness from each subunit was replicated and summed over the structure prior to averaging. To understand the role of cofactors for communicating allostery, we extended this analysis with a measure of signaling efficiency. Here, efficiency is characterized by *current flow closeness centrality* [18] (see Supplementary Material). We thus used current flow betweenness and closeness centrality to extract allosterically important protein residues, and to reveal the allosteric role of lipids and ligands. To avoid confusion with ion channel currents, we will use *information flow* to denote current flow betweenness and *information flow closeness* to denote current flow closeness.

We studied three well-known systems; the voltage-gated ion channel KCNQ1, the GPCR *β*2 adrenergic receptor, and the pH-gated ion channel KcsA. Technical details about simulation setup and protocols are described in Supplementary Material. The structures were visualized with VMD [35].

## IV. RESULTS AND DISCUSSION

### A. KCNQ1 exploits the membrane for communicating voltage sensitivity to the pore

KCNQ1 is a voltage-gated potassium channel critical for action potential repolarization in the heart. KCNQ1 contains the common elements of a homotetrameric voltage-gated ion channel; each subunit consists of six transmembrane helices [36]. The first four helices form the voltage-sensing domain (VSD), while the last two make up the pore domain, Fig. 2A. The Ca^2+^-sensor calmodulin (CaM) is required for KCNQ1 channel assembly [37, 38], and binds to the KCNQ1 C-terminal domain (CTD) [36, 39]. Specific to KCNQ1 is that PIP_2_ is required to couple VSD activation to pore opening [40, 41]. The two cryo-EM structures of human KCNQ1 [39] revealed “bent” CTD helices in the absence of PIP_2_, and “straight” CTD helices in the presence of PIP_2_, Fig. S2. We call these states *CTD-bent* and *CTD-straight*, respectively. The CTD-bent state allows interactions between CaM and the VSD, which are disrupted in the CTD-straight channel. Recent studies showed that both states are present in the channel activation pathway [21].

**FIG. 2.**
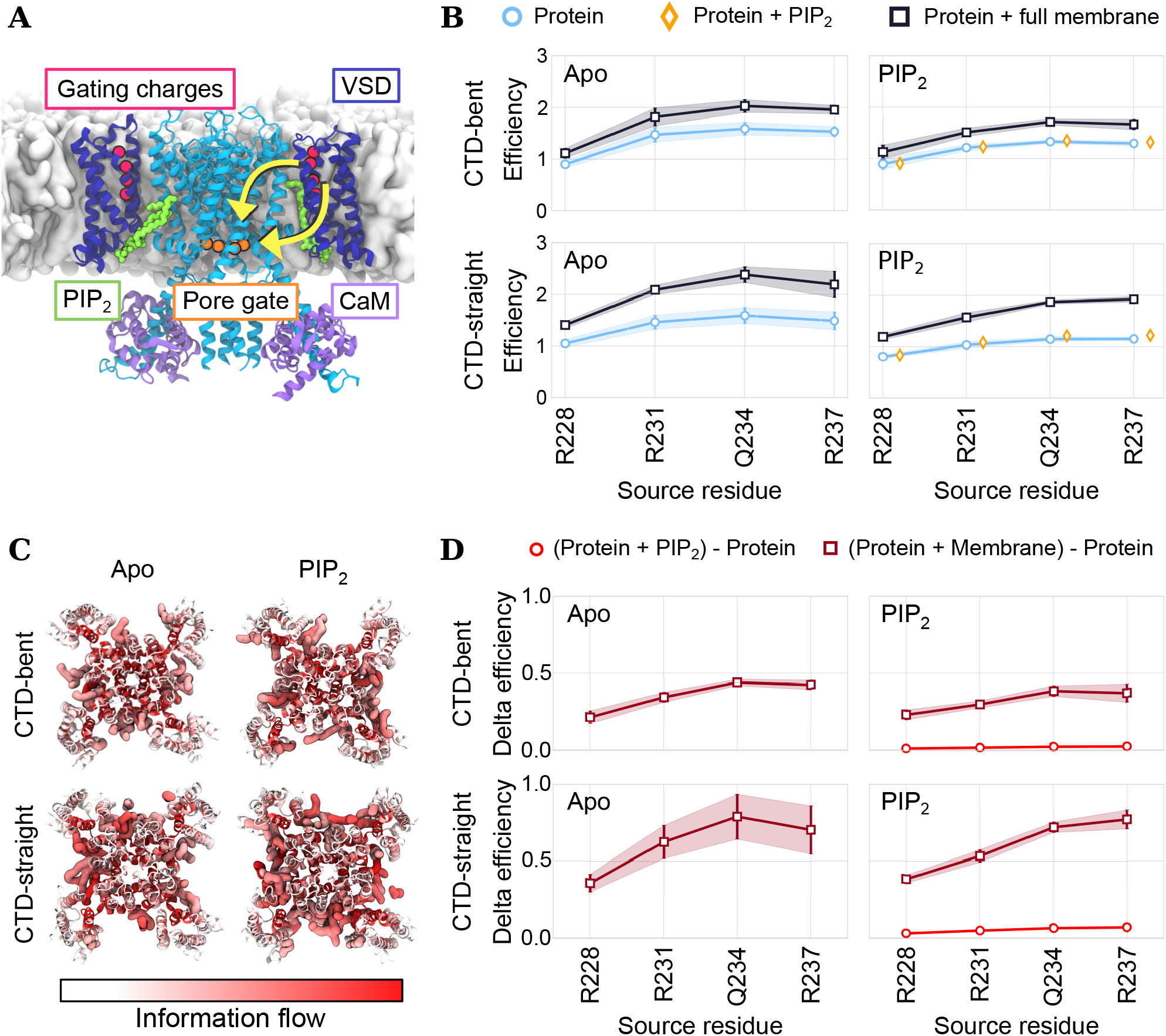
**(A)** Structure of CTD-straight KCNQ1 with important structural features, and sources and sinks. The yellow arrows show cartoon pathways, highlighting the direction between the source and sink residues. **(B)** State- and PIP_2_-dependent allosteric efficiency (information flow closeness) measured from the source residues listed on the x-axis to the sink residue S349. Blue: protein, yellow: protein+bound PIP_2_, and black: protein+full membrane. Left column: apo, and right: PIP_2_-bound. Upper row: CTD-bent state, and bottom row: CTD-straight. **(C)** Average information flow through each protein residue and lipid interactor projected onto the first trajectory frame (top view). Red signifies larger information flow values. Lipid interactors (information flow > 0.008) are shown in surface representation. **(D)** Difference in allosteric efficiency of networks including different components of the systems. Dark red: comparing protein/membrane networks to protein networks, and red: comparing protein/PIP_2_ networks to protein networks. Averages and standard errors (SEM) are computed across subunits.

A thorough investigation of the KCNQ1 protein information flow from the voltage-sensor domain to the pore, validated by electrophysiology, is presented elsewhere [21]. Here, on the other hand, we studied the allosteric roles of PIP_2_ and the membrane to allow communication between the VSD and pore. Efficiency of communication was measured in terms of information flow closeness using the KCNQ1 gating charges (R1, R2, Q3. R4) as sources and the pore residue S349 as sink. The allosteric contribution of each system component was inferred by performing the efficiency analysis on the networks including: 1) the protein, 2) the protein and PIP_2_, and 3) the protein and the full membrane. Fig. 2B shows the allosteric efficiency when including PIP_2_ and the lipid membrane in the network. The upper row displays the results obtained from the CTD-bent simulations, while the lower row shows the corresponding results from the CTD-straight simulations. The left and right columns correspond to PIP_2_ being absent or present in the simulations, respectively. Regardless of PIP_2_-bound state, the membrane increases the allosteric efficiency, indicating that the membrane is involved in communicating voltage sensitivity to the pore, Fig. 2B. Fig. 2C depicts a top-view of the information flow through each residue projected onto the CTD-bent (upper) and CTD-straight (lower) structures. The lipid interactors with the highest information flow are shown in surface representation, indicating their location with respect to the protein. KCNQ1 exploits the membrane lipids tucked in the cavities between VSDs and the pore to communicate allostery, Fig. 2C. Fig. 2D shows the difference in efficiency between networks which include the additional interactors compared to the isolated protein. PIP_2_ appears to not significantly increase signaling efficiency alone. The effect of tightly bound ligands may thus already be encoded in the isolated protein network. Moreover, the CTD-straight channel uses the membrane to a larger extent than the CTD-bent, while PIP_2_ appears to not alter the involvement of the membrane. Previous studies suggested that state-dependent PIP_2_-binding is important for stabilizing different channel conformations [41]. Accordingly, our results suggest that the role of the bound PIP_2_ is not to explicitly couple the VSD to the pore via the membrane, but to induce the large rearrangements in the CTD related to pore opening.

### B. Agonist ligand-binding decreases the membrane allosteric role in the inactive *β*2 adrenergic receptor

The *β*2 adrenergic receptor (*β*2AR) is a G-protein coupled receptor (GPCR) and a common target for asthma medication. It sustains communication through the cell membrane by recognizing and binding ligands. The binding of agonist ligands in the receptor extracellular domain induces structural rearrangements throughout the protein. The induced state, known as the active state, allows for intracellular G-protein association, which triggers cellular downstream responses [6, 42, 43]. As such, understanding the molecular mechanisms of agonist-imposed allosteric signaling between the two binding sites may be an essential element to facilitate development of novel drugs.

Fig. 3A shows the structural elements of the *β*2AR with the bound agonist ligand BI-167107 and G-protein mimetic nanobody (Nb) [44]. We applied the information flow analysis to the agonist bound as well as ligand-free (apo) *β*2AR in the active/Nb-bound and inactive/Nb-free states, respectively. To investigate the allosteric roles of the agonist and membrane, we performed the analysis on networks including different system components: the protein, the protein and agonist, as well as the protein, agonist and membrane. Source nodes were defined as residues in the orthosteric site which are known to interact with agonists, Fig. 3A-B [45]. The sink nodes were taken as residues in contact with the G protein in the active state structure 3SN6 [46], Fig. 3A and Table S1.

**FIG. 3.**
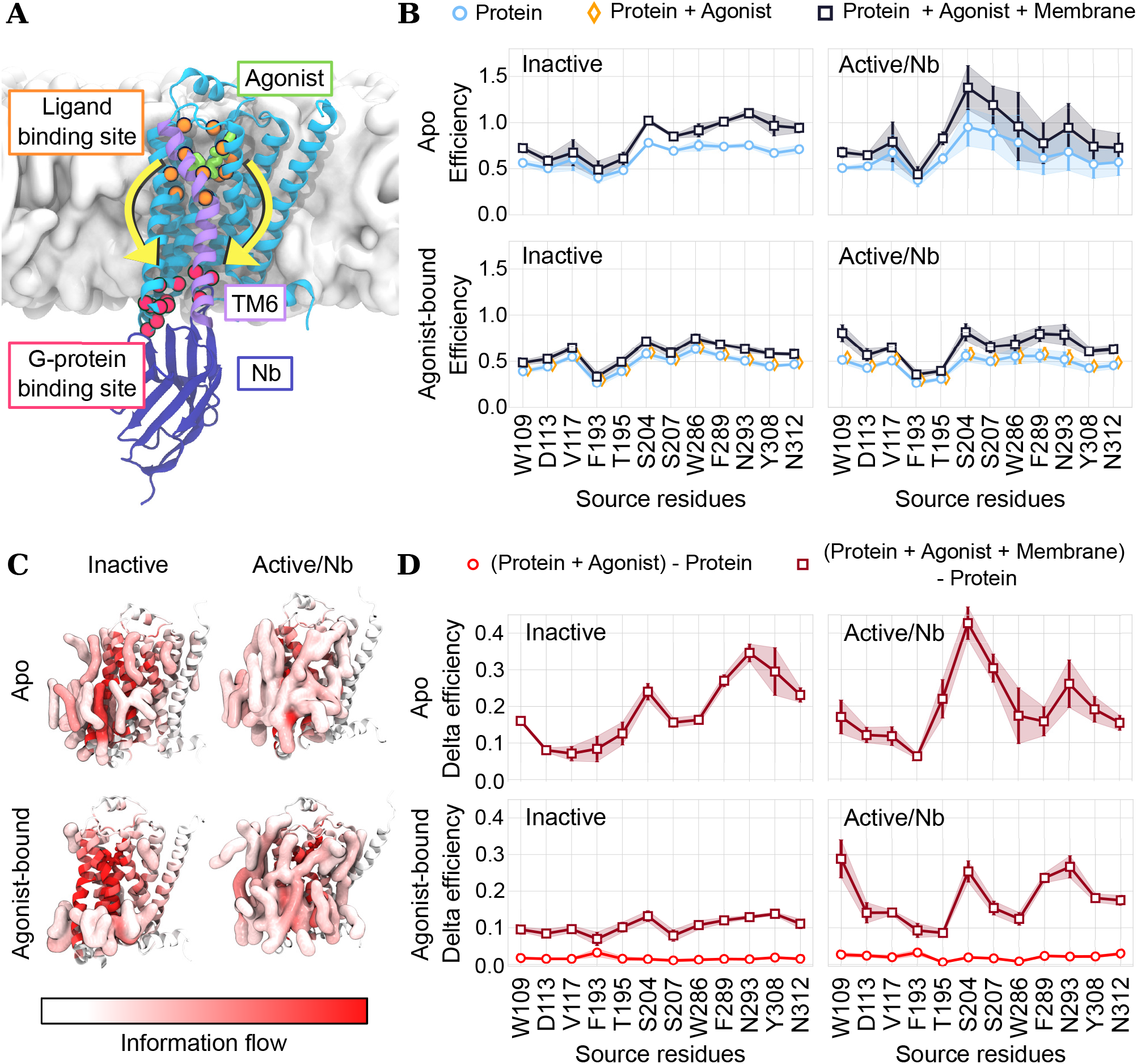
**(A)** Structure of active/Nb-bound *β*2AR showing important structural features, and sources and sinks. The yellow arrows show cartoon pathways, highlighting the direction between the source and sink residues. **(B)** State- and ligand-dependent allosteric efficiency (blue: protein, yellow: protein + agonist, black: protein + agonist + full membrane). Left column: inactive state, and right: active/Nb-bound state. Upper row: apo *β*2AR, bottom row: agonist-bound *β*2AR. **(C)** Average information flow through each protein residue and lipid interactors (information flow > 0.012) projected onto the first trajectory frame. Red signifies larger information flow values. **(D)** Difference in allosteric efficiency of networks built by including different components of the *β*2AR systems. Dark red: comparing protein/agonist/membrane networks to protein networks, and red: comparing protein/agonist networks to protein networks. Averages and SEM are obtained by splitting the simulations into three parts.

The apo inactive *β*2AR exploits the membrane to transfer allostery between the ligand and G-protein binding sites, as shown by an increased signaling efficiency after including the membrane in the network Fig. 3B. Specifically, it uses the closest layer of the lipid membrane, Fig. 3C. The agonist-bound inactive receptor, on the other hand, demonstrates a significantly decreased allosteric involvement of the membrane relative to the apo receptor, Fig. 3B-D. This is shown by the smaller increase of signaling efficiency when including the membrane, Fig. 3B,D, as well as a lower amount of information flow through lipids, Fig. 3C. This suggests that the bound ligand decouples the allosteric pathways from the membrane in the inactive state. Conversely, the membrane increases allosteric efficiency of the active/Nb-bound receptor, in both apo and agonist-bound systems, Fig. 3B-D. Thus, an intracellular binding partner may again allosterically couple *β*2AR to the surrounding membrane in its active state.

To understand the allosteric details related to exploiting the membrane, we plotted the information flow along protein residue sequence and contrasted the profiles obtained from the apo (black) and agonist-bound (blue) simulations, Fig. 4A. The profiles show a large allosteric involvement of TM3 and TM5-7. These helices are important for activation and interact with both binding partners [46–49]. An interesting feature related to ligand binding emerged; namely an increased involvement of TM1, in particular of the residues I43, V44, I47 and N51, Fig. 4B. N51 is highly conserved across all class A GPCRs [50] and supports a stabilizing water network in the receptor’s inactive state [51]. Notably, the inactive agonist-bound receptor also exhibited more information flow at TM3, compared to the apo inactive receptor. The difference between agonist-bound and apo information flow profiles is smaller in the active/Nb state than in the inactive state. Together with the results of Fig. 3D, this indicates that an intracellular binding partner overshadows the ligand-induced redirection of allosteric pathways within the protein.

**FIG. 4.**
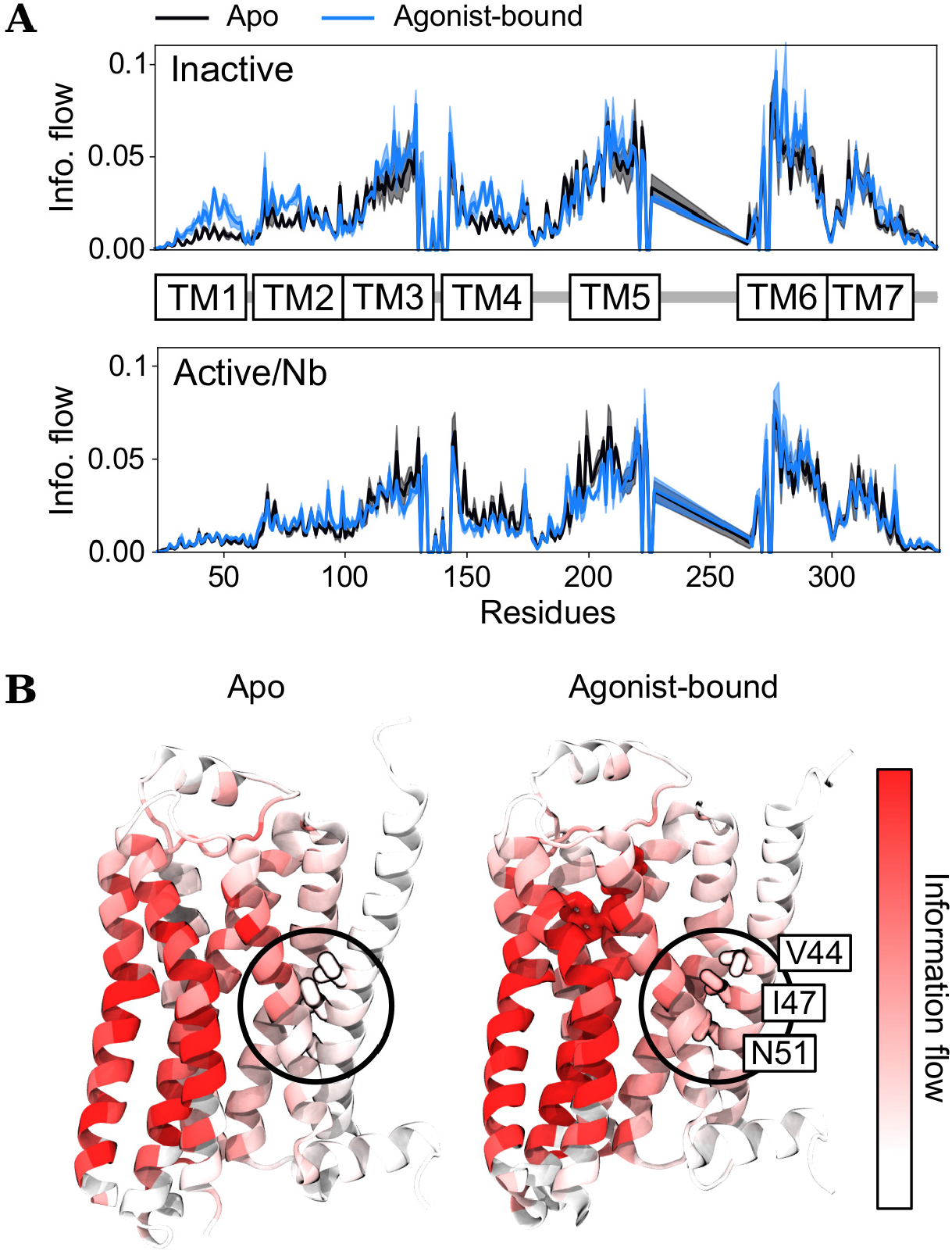
**(A)** Average and standard deviation of protein information flow plotted along residue sequence of apo (black) and agonist-bound (blue) *β*2AR in inactive (upper) and active/Nb (lower) states. **(B)** information flow projected onto apo (left) and agonist-bound (right) inactive structures. The black circle shows a region (encompassing TM1, TM2 and TM7) which becomes more involved upon agonist-binding. Averages and standard deviations are obtained by splitting the simulations into three parts.

### C. Binding of DOPG lipid changes the KcsA allosteric signaling at the open state

KcsA is a bacterial pH-gated and homotetrameric potassium channel [52]. The simplicity and high conservation of structural features make the channel a well-suited model for complex eukaryotic potassium channels [53]. Mechanisms related to its gating and lipid regulation are therefore widely studied [53–56]. Two structural components are important for conduction of potassium ions in KcsA: the *selectivity filter* (SF) and the *inner gate*, Fig. 5A. The SF, which selectively coordinates potassium ions, is located at the extracellular side of the protein. The sequence of SF residues is highly conserved across potassium channels [57]. The inner gate, on the other hand, is located on the intracellular side of the protein. It is formed by the crossing of the second transmembrane helix (TM2) from each subunit [53]. A pH drop in the cell causes the inner gate to open, yielding the KcsA open state channel. Within milliseconds after opening, the selectivity filter collapses, which hinders ion permeation. This process, known as *inactivation*, is crucial to control electrical signals and membrane potentials in excitable cells [57]. The constriction of the SF, which is responsible for inactivation, is thus allosterically triggered by a structural change at the inner gate [56].

**FIG. 5.**
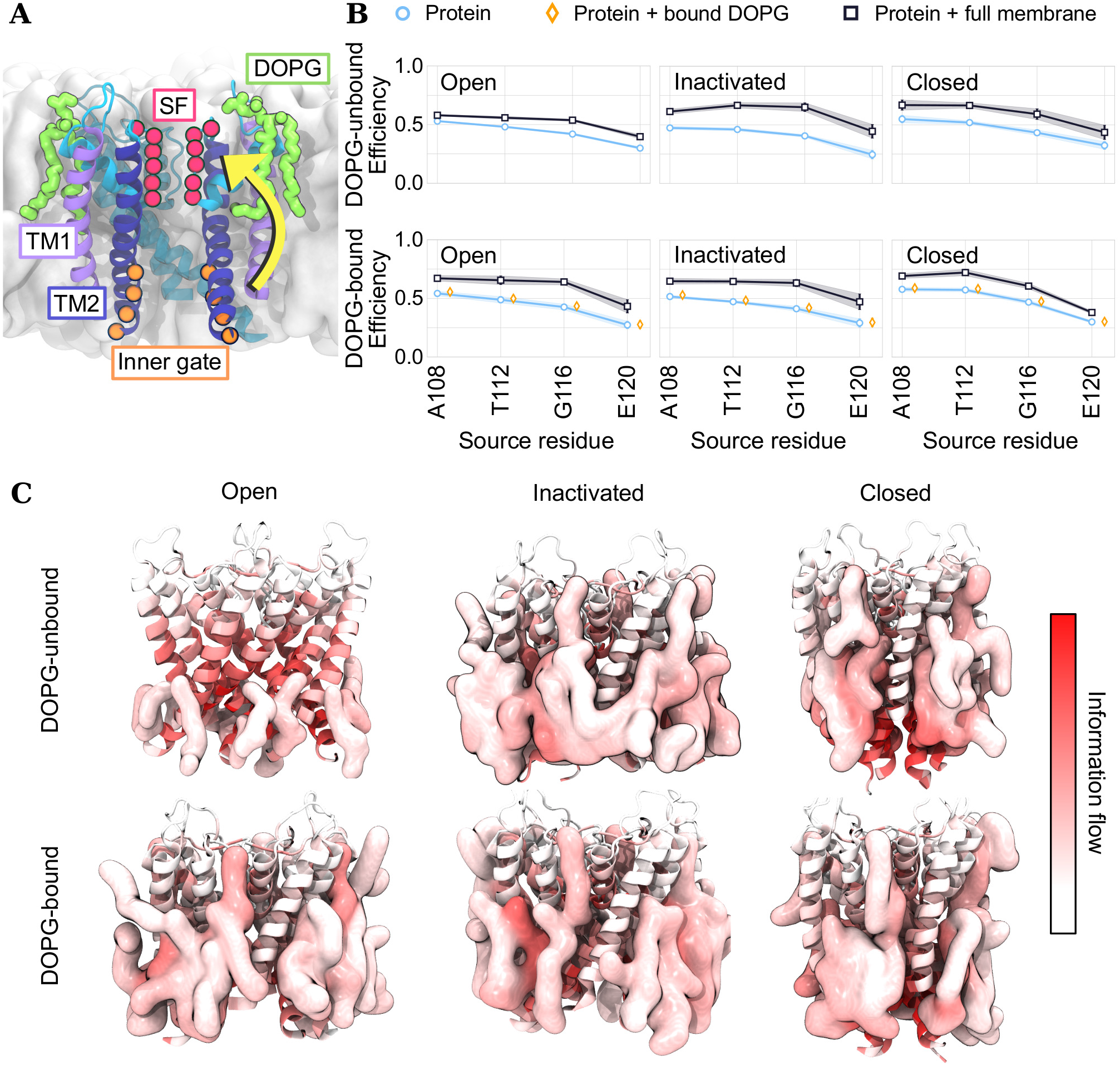
**(A)** Structure of open/DOPG-bound KcsA showing important structural features, and sources and sinks. The subunit in front of the channel pore is omitted for clarity. The yellow arrow shows a cartoon pathway, highlighting the direction between the source and sink residues. **(B)** State- and DOPG-dependent allosteric efficiency (blue: protein, yellow: protein + bound DOPG, black: protein + full membrane). Left column: open state, middle: inactivated state, and right: closed state. Upper row: DOPG-unbound channels, and bottom row: DOPG-bound channels. **(C)** Average information flow through each protein residue and lipid interactor (information flow > 0.012) projected onto the first trajectory frame. Red signifies larger information flow values. Averages and SEMs are computed across subunits.

Lipids are important for KcsA modulation. The membrane thickness, for example, affects the open state probability of the E71A mutant [58]. In addition to this, experiments indicate that inactivation is facilitated by the binding of an anionic lipid to each intersubunit cavity, Fig. 5A [55]. These results are obtained by mutating residues at the lipid binding sites. Such mutations may, however, yield unknown structural effects. In contrast, the approach suggested here allows for accessing the molecular details in the wild-type channel, given the general limitations of molecular modeling.

We investigated the allosteric role of a bound DOPG molecule and of the lipid membrane in the open, inactivated and closed states using the inner gate residues A108, T112, G116 and E120 as sources and SF residues T75 to G79 as sinks, Fig. 5A. Fig. 5B shows the state-dependent allosteric efficiency of each system component; the protein, the protein and the bound DOPG, and the protein and the full membrane. The DOPG-unbound open channel appears to allosterically rely on the membrane slightly less than the DOPG-unbound inactivated and closed channels, Fig. 5B. The DOPG-bound open channel uses lipids located at the intracellular side of the protein to transmit allosteric signals, Fig. 5C. The inactivated and closed channels, on the other hand, exploit lipids from the intersubunit surface at the upper leaflet all the way down to the lower leaflet and the inner gate, Fig. 5C. Interestingly, DOPG binding appears to specifically increase the allosteric role of the lipid membrane in the open state channel, such that the bound lipid connects the upper and lower leaflet lipids, Fig. 5B,C.

To further investigate KcsA allostery related to the open state, we compared information flow profiles between different states of DOPG-unbound and DOPG-bound channels, Fig. 6A, S3. This revealed differences at the central residues of TM2 helix, and particularly at the TM1 helix, Fig. 6A,B, S3. The allosteric importance of TM1 (W26-L40) is increased in the DOPG-unbound open state channel. L40 is specifically important for the coupling between inner gate and SF at the open state [59]. The increased importance of this region in the DOPG-unbound state may be a consequence of the decreased allosteric involvement of the membrane. We hypothesize that the bound DOPG promotes inactivation by coupling the protein to the membrane, thereby altering the allosteric pathways within the protein.

**FIG. 6.**
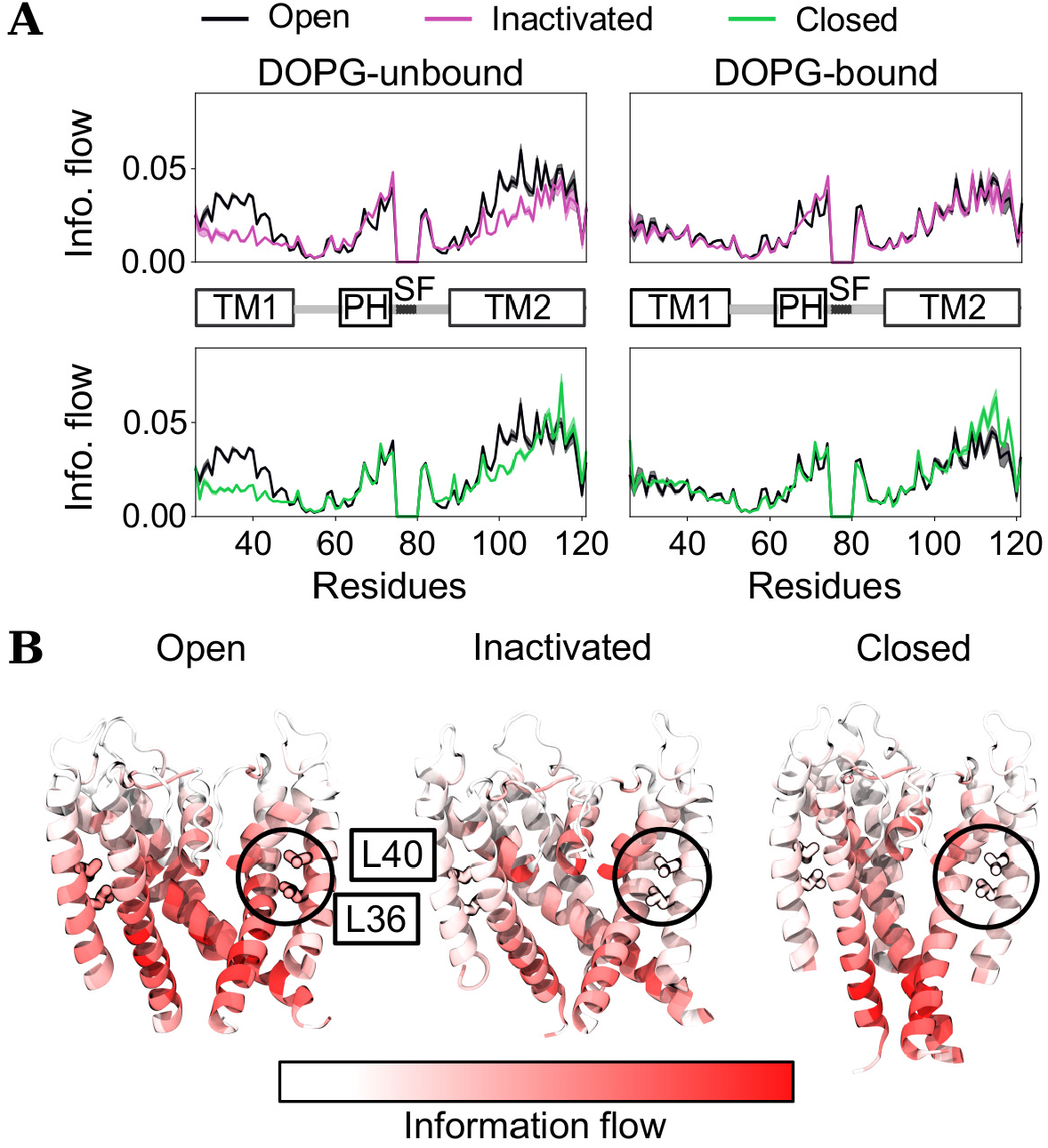
**(A)** Average and standard deviation of protein information flow plotted along residue sequence of open (black), inactivated (purple) and closed (green) states with (right) and without (left) bound DOPG. KcsA structural features are labeled along sequence: TM1, pore helix (PH), selectivity filter (SF), TM2. **(B)** information flow projected onto the DOPG-unbound protein structures. The circle shows the region on TM1 and TM2 which is more allosterically involved in the open/DOPG-unbound channel. The subunit in front of the channel pore is omitted for clarity. Averages and standard deviations are computed across subunits.

## V. CONCLUSIONS

We presented a framework which extends ordinary residue interaction networks to include cofactors such as lipids and ligands. By solving a linear assignment problem, and thus optimally assigning interactors to nodes in each frame, the problem of cofactor redundancy is avoided. We applied this framework to obtain networks of three different types of membrane proteins and their cofactors. The network analysis revealed interesting features of protein allostery modulation by lipids and ligands. Importantly, we showed that the lipid membrane plays an integral role in the state-dependent allosteric communication of these proteins. The strong involvement of the membrane in KCNQ1 allostery may be attributed to the nonconvex structural topology of KCNQ1. The *β*2AR and KcsA datasets specifically demonstrated that subtle redirections of allosteric pathways induced by a modulator may change the membrane involvement. Network analysis including cofactors may thus serve to reveal intricate allosteric mechanisms of membrane proteins.

## AUTHOR CONTRIBUTIONS

L.D. initiated and coordinated the project. L.D. and A.M.W. defined the project scope. A.M.W. developed and implemented the framework, and analyzed the datasets. O.F., S.P.C. and A.M.W. performed simulations of the *β*2 adrenergic receptor, KcsA and KCNQ1, respectively. A.M.W. prepared the figures and the first manuscript draft. All authors participated in interpreting the results and writing the paper.

## SUPPLEMENTARY MATERIAL

Supplementary material includes additional Methods section, Figs. S1-S3, and Table S1. The Methods section describes MD simulation parameters and information flow (current flow) calculations.

## AIP PUBLISHING DATA SHARING POLICY

The data that support the findings of this study are openly available in the open science framework repository https://osf.io/2c6zd/, [60]. Code and tutorial for building and analyzing networks are available at https://www.github.com/delemottelab/allosteric-pathways [20].

## ACKNOWLEDGMENTS

This work was supported by grants from the Science for Life Laboratory, the Göran Gustafsson Foundation and the Swedish research council (VR-2018-04905). The simulations were performed on resources provided by the Swedish National Infrastructure for Computing (SNIC) at PDC Centre for High Performance Computing (PDC-HPC). The authors thank Christian Blau for insightful discussions during method development.

