## Supplementary text and figures for "Network analysis reveals how lipids and other cofactors influence membrane protein allostery"

### I. SUPPLEMENTARY METHODS

#### A. Current flow betweenness and current flow closeness centrality

Current flow is obtained from the network Laplacian, $L = D - A$, where $A$ is the adjacency matrix and $D$ is the diagonal matrix obtained by summing the rows of $A$. The rows and columns corresponding to sink nodes are removed from the Laplacian and the matrix is inverted. By adding zero rows and columns at the sink nodes again, we obtain the inverse reduced Laplacian, $\tilde{L}^{-1}$. The potentials are given by

$$p(s) = \tilde{L}^{-1}b, \tag{1}$$

where $b$ is the supply vector which corresponds to one unit of current entering at the source node ($b_s = 1$) that will exit at sink nodes. The current flow betweenness, $\tau_i$, of node $i$ from all source nodes to sink nodes is obtained from the average over sources,

$$\tau_i = \frac{1}{|S_0|}\sum_{s\in S_0} f_i(s) \tag{2}$$

where

$$f_i(s) = \begin{cases} 0 & \text{if } i \in S_1 \text{ or } i = s \\ \frac{1}{2}\sum\limits_{j} |p_i(s) - p_j(s)|A_{ij} & \text{otherwise} \end{cases} \tag{3}$$

By setting the sink and source node current flows to zero, they are removed from the analysis.

Current flow closeness centrality estimates efficiency as the average inverse potential difference between different source and sink pairs [1]. Current flow closeness between a specified source $s$ and the sinks nodes is given by

$$\eta_s = \frac{1}{\frac{1}{|S_0|}\sum\limits_{t\in S_1} |p_s(s) - p_t(s)|}, \quad \forall s \in S_0. \tag{4}$$

#### B. MD simulation details

##### 1. KCNQ1

We built four systems with human KCNQ1 using the cryo-EM structures obtained in the presence and absence of the bound $PIP_2$ (PDB IDs: 6UZZ, 6V01 without KCNE3) [2] using CHARMM-GUI [3, 4]. The system preparation was done following prior studies of KCNQ1 [5, 6]. In short, missing loops were modeled using modeller and the DOPE score [7] and the protein was embedded in a POPC membrane bilayer [8] of $\sim 340$ molecules, then solvated in a 0.1M concentration of neutralizing KCl and TIP3P [9] water molecules. Equilibration was carried out using the 6-step Charmm-gui procedure [4], extending the sixth step to 3 ns.

The systems were simulated with Gromacs 2018 [10] using the CHARMM36 force field [11], resulting in three replicas with a total simulation time of $\sim 1\mu$s per system. The Nose-Hover thermostat [12] (T = 300K, $\tau = 1$ps) and semi-isotropic Parrinello-Rahman barostat (P = 1bar, $\tau = 5$ps) [13] were used. Bonds involving H-atoms were constrained with LINCS [14]. Long-ranged electrostatics were calculated using PME (rcoulomb = 1.2nm fourierspacing = 0.15nm$^{-1}$ and order = 4) [15]. Van der Waals interactions were force-switched to 0 for distances between 1.0 nm and 1.2 nm. Unless otherwise stated, the following simulation systems will use the same parameters.

#### 2. *The $\beta$2 adrenergic receptor*

The $\beta$2 adrenergic receptor systems were built from a nanobody-bound active-state structure (PDB ID: 3P0G) [16] in CHARMM-GUI [3, 4] with the CHARMM36m force field [17]. Although the nanobody-bound $\beta$2AR is not a native state, nanobodies are often used as G-protein proxies to lock GPCRs in active states which are highly similar to their native counterparts [18]. Following the protocol of a prior study [19], we reversed the N187E in the crystallized structure, protonated E122(3.41), and protonated the two histidines H172(4.64) and H178(4.70) at their epsilon positions. The receptor was embedded in a POPC membrane bilayer [8] of 180 or 190 molecules depending on whether the nanobody was included or not, then solvated in a 0.15M concentration of neutralizing sodium and chloride ions with 79 or 120 TIP3P [9] water molecules per lipid molecule depending on presence of nanobody.

An adaptive sampling protocol was applied to sample the most stabilized state that is kinetically accessible from the starting structure without applying an artificial force on the system [20]. Briefly, 24 simulation replicas of 7.5 ns were initially launched. The replicas' center point, c, in CV space was taken as the mean of the trajectories' endpoint coordinates, $x_i$. The CVs to describe and distinguish conformational states are obtained from 41 weighted inverse inter-residue distances identified as important using the software demystifying [20, 21]. The center point and average distance, $d$, from the endpoints to the center were used to assign a weight to each replica, $i$,

$$w_i(x) = \exp\left[-\frac{|x_i - c|^2}{d^2}\right]. \tag{5}$$

New trajectories were iteratively seeded by extending

$$n_i \propto \frac{w_i}{\sum_j w_j} \tag{6}$$

copies of each replica, keeping the total number of trajectories fixed to 24. This approach favors replicas close to the center, leaving the ensemble of trajectories to diffuse around a single equilibrated state for a total of 1.4$\mu$s per system.

#### 3. *KcsA*

The systems were built from the closed, open and inactivated structures (PDB ID:5VKH, 5VK6 and 5VKE) [22, 23] in CHARMM-GUI [3, 4] with the CHARMM36 force field [11]. We did not include the protein's C-terminus and pH sensor. Although we note that the missing pH sensor may influence the stability of conformational states, we chose to keep the protein chain lengths consistent across states. Given available KcsA structures, we therefore kept residues 26-121. Mutations were reverted with PyMOL. E71, E118 and E120 were protonated; the first to form the important interaction with D80, and the others in order to simulate channel activation conditions at pH=4. The N- and C-termini were capped by acetylation (NME) and methylation (ACE). The selectivity filter was fully loaded with potassium ions. The protein was embedded in a DOPE/DOPG 3:1 membrane bilayer of  150 molecules, then solvated in a 0.1M concentration of neutralizing KCl and NaCl with 75 TIP3P  [9] rigid water molecules per lipid molecule. Each state was simulated with and without co-purifying lipids (DOPG-bound/DOPG-unbound), as originally identified in X-ray structures [22, 24].

1$\mu$s long MD simulations were performed with gromacs2019 [10]. The v-rescale thermostat [25] (T = 290K $\tau$ = 1ps) and semi-isotropic Parrinello-Rahman barostat (P = 1bar, $\tau$ = 5ps) [13] were used. Parts of the simulations where a water molecule enters the selectivity filter to cause backbone flips similar to pre-inactivation were omitted from the analysis.

### II. SUPPLEMENTARY FIGURES

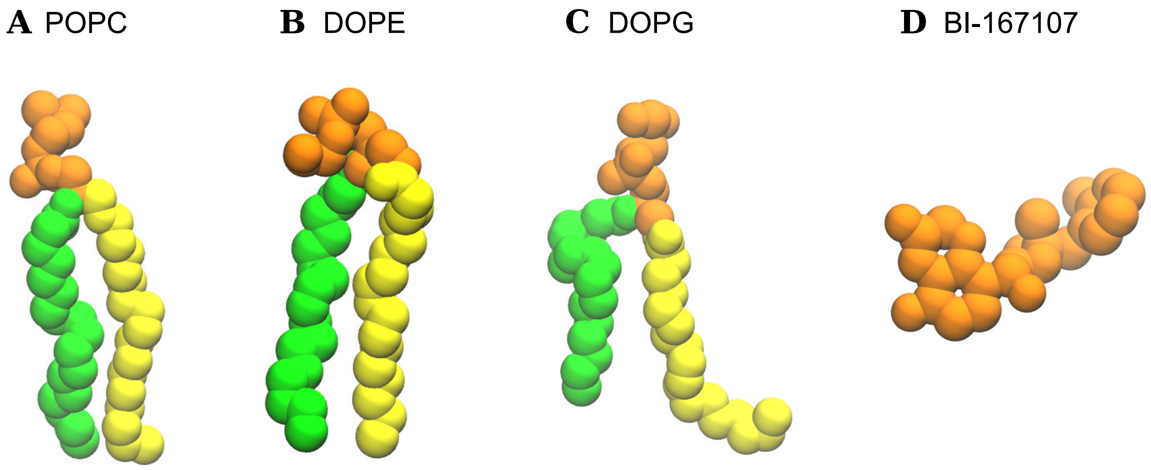


FIG. S1. The grouping of a) POPC, b) DOPE, c) DOPG lipids, and d) the BI-167107 ligand into the domains (interactors) used to define nodes in the networks. The atoms involved in different interactors are shown in different colors.

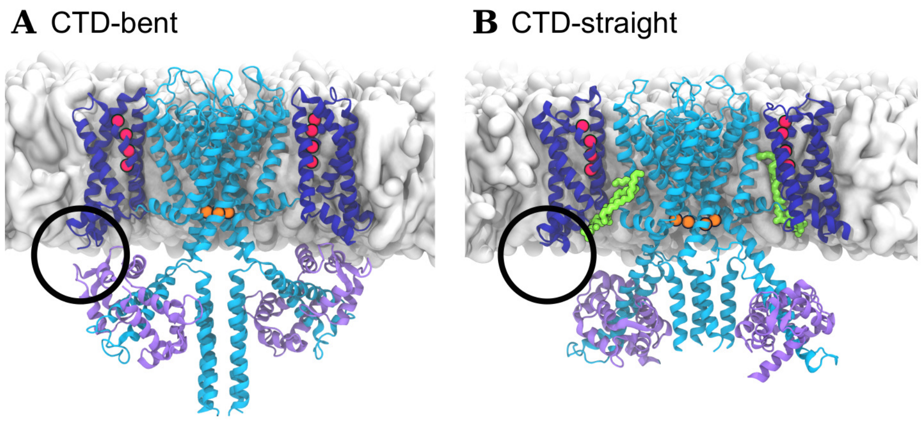


FIG. S2. The KCNQ1 in a) CTD-bent and b) CTD-straight states. The black circle highlights the presence and absence of interactions between CaM and the KCNQ1 voltage sensing domain.

TABLE S1. Sink residues used for analysis of the $\beta$2AR systems.

| |
|---|
| R131 |
| A134 |
| I135 |
| T136 |
| P138 |
| F139 |
| Y141 |
| Q142 |
| S143 |
| V222 |
| E225 |
| A226 |
| R228 |
| Q229 |
| L230 |
| K232 |
| I233 |
| A271 |
| T274 |
| L275 |

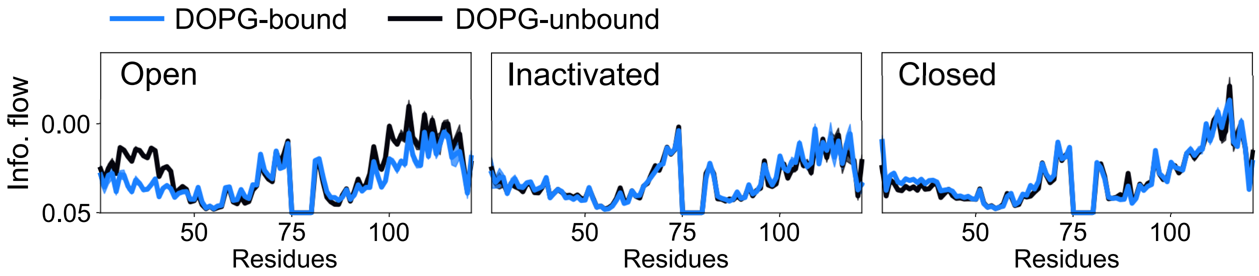


FIG. S3. Current flow profiles contrasting DOPG-bound to DOPG-unbound KcsA channels.

[17] J. Huang, S. Rauscher, G. Nawrocki, T. Ran, M. Feig, B. L. de Groot, H. Grubmüller, and A. D.

MacKerell, Nature Methods **14**, 71 (2017).
